# Temporal uncertainty enhances suppression of neural responses to predictable visual stimuli

**DOI:** 10.1101/2020.08.26.265884

**Authors:** Sanjeev Nara, Mikel Lizarazu, Craig G Richter, Diana C Dima, Radoslaw M Cichy, Mathieu Bourguignon, Nicola Molinaro

## Abstract

Predictive processing has been proposed as a fundamental cognitive mechanism to account for how the brain interacts with the external environment via its sensory systems. In vision, contextual information triggers predictions about the content (“*what*”) of environmental stimuli to update an internal generative model of the surrounding world. However, visual information dynamically changes across time, and temporal predictability (“*when*”) may influence the impact of internal predictions on visual processing. In this magnetoencephalography (MEG) study, we investigated how processing feature specific information (“*what*”) is affected by temporal predictability (“*when*”). In line with previous findings, we observed suppression of evoked neural responses in the visual cortex for predictable stimuli. Interestingly, we found that temporal uncertainty increased expectation suppression. This suggests that in temporally uncertain scenarios the neurocognitive system invests less resources in integrating bottom-up information. Multivariate pattern analysis showed that predictable visual features could be decoded from neural responses. Temporal uncertainty did not affect decoding accuracy for early visual responses, with the feature specificity of early visual neural activity preserved across conditions. However, decoding accuracy was less sustained over time for temporally jittered than for isochronous predictable visual stimuli. These findings converge to suggest that the cognitive system processes visual features of temporally predictable stimuli in higher detail, while processing temporally uncertain stimuli may rely more heavily on abstract internal expectations.

## Introduction

Our interaction with the external environment is largely shaped by our internal expectations (Clark, 2013; Mechelli et al., 2004; Mumford, 1992). In the primary visual cortex, percepts are decomposed into their fundamental features (such as edges, orientations, colours, shapes) and dissociable correlates of such representational properties can be decoded from neural signals (Carlson et al., 2019; Pantazis et al., 2018). These features are the building blocks that determine the perceptual content – “what” we perceive a stimulus to be. Perception of content is largely modulated by higher level processes. Predictive processing theories propose that an internal mental model of the surrounding environment is used to generate inferences about the external causes of the environmental energy impacting our senses (Spratling, 2017). Supporting this view, responses in the visual cortex to predictable visual stimuli have reduced amplitudes and are processed at shorter latencies than unpredictable stimuli (Hogendoorn and Burkitt, 2018). It is worth underscoring that perception is spatio-temporal in nature since, in real life situations, environmental stimuli are temporally dynamic. Suppose you see a car coming toward you on the road. Even after determining all its visual features (“what” you see, e.g., *a pink Cadillac*), you will still need to estimate *when* you and the car would intersect to avoid a collision. While certain stimuli are temporally regular (and hence temporally predictable), the temporal uncertainty of the natural environment is very high.

Studies on visual perception have paid relatively less attention to temporal (*when*) than to content (*what*) predictability (Demarchi et al., 2019; Kok et al., 2017). To date, the role of timing in predictive processing in vision has mainly been studied in relation to the perception of objects in motion. The visual system requires a certain amount of time to process incoming sensory information (Blom et al., 2020; Maunsell and Gibson, 1992). Yet, the neurocognitive system must rapidly extrapolate the trajectory of moving objects to expedite actions. Predictive processing accounts propose a compensatory mechanism to support such extrapolations: the system generates predictions about incoming stimuli, which trigger visual responses before the temporal onset of the actual stimulation. Research on the perception of moving objects thus underscores that *what* and *when* stimulus properties are strongly interwoven and shape human perception. Here we aim to evaluate how predictive processing of “*what* properties” is affected by the manipulation of “*when* properties”. Specifically, we test the hypothesis that temporal uncertainty enhances predictive processing of stimulus content. We base this hypothesis on the theoretical claim that sensory systems require internal predictions in order to deal with uncertainty in the external environment; increased uncertainty leads to increased reliance on predictive processing (Clark, 2013).

In the present MEG study, participants viewed four consecutive Gabor patches (henceforth *entrainers*), then had to make a spatial frequency judgment on a. fifth *target* Gabor. Entrainers followed either a predictable or unpredictable sequence of orientations and had either isochronous or jittered onset times. We planned four main analyses. Firstly, we expected neural responses to entrainers to gradually decrease in amplitude across four-element sequences when orientations were predictable due to “expectation suppression” (Grill-Spector et al.,2006). Secondly, we hypothesized that if the cognitive system handles temporal uncertainty by increasing its reliance on internal predictions, expectation suppression should be larger for onset-jittered compared to onset-isochronous entrainers. Thirdly, we expected multivariate pattern analysis (MVPA; Pantazis et al., 2018; King et al., 2016; Cichy et al., 2014: Grootswagers et al., 2017) to reveal increasingly high decoding accuracy of entrainer orientation for predictable entrainers, reflecting the brain’s incrementally higher reliance on predictions for successive Gabor orientations. Finally, we planned to analyze the time course of entrainer orientation classification results to understand how (both *what* and *when*) predictability affects the stimulus specificity of the neural responses.

## Methods

### Participants

From the initial set of twenty participants, we included data from sixteen participants (7 females; age range: 19–31; M = 24.8; SD = 3.6) in our analyses. Two participants were excluded from the study as they did not complete the whole experiment and two more were excluded from the study due to excessive motion artifacts in their data. The ethical committee and the scientific committee of the Basque Center on Cognition, Brain and Language (BCBL) approved the experiment (following the principles of the Declaration of Helsinki). Participants gave written informed consent and were financially compensated. The participants were recruited from the BCBL Participa website (https://www.bcbl.eu/participa/). Participants did not present any neurological or psychological disorders, and had normal or corrected to normal vision.

### Experimental procedure

A series of Gabor patches with variable orientations and spatial frequencies (measured in cycles per degree [CPD] of visual angle) was presented. Stimuli were back-projected on a screen placed 60 cm from each participant’s nasion. The Gabors were presented in the centre of the screen on a gray background, covering the central two degrees of the visual field. Each trial began with a fixation cross (black color) followed by four sequential Gabor patches (entrainers), each presented for 200 ms followed by an interstimulus interval showing an empty grey screen. After a longer interstimulus interval, a fifth Gabor (target) was presented for 200 ms. The entrainers had an intermediate spatial frequency (40 CPD), while the target could have either a higher (60 CPD) or lower (20 CPD) spatial frequency. Participants were required to indicate if the target had a higher or lower spatial frequency than the entrainers using a button press.

Four properties of these sequences were experimentally manipulated (Figure 1): a) the orientation of the target was either horizontal or vertical; b) the spatial frequency of the target was either higher or lower than the spatial frequency of the entrainers; c) the orientation of the target was either predictable based on the orientations of previous entrainers (i.e., clockwise or counter-clockwise rotations of either 15 or 30 degrees; e.g., entrainers of 30, 45, 60, 75 and a target of 90 degrees) or unpredictable (a random selection from a set of 15 or 30 degree rotations; e.g., 30, 75, 60, 45 and a target of 90 degrees); d) the timing of the interstimulus intervals (blank grey screens) between the four entrainers and between the last entrainer (entrainer 4) and the target was either predictable (i.e., fixed interstimulus intervals of 200 ms between entrainers and 600 ms between entrainer 4 and the target) or unpredictable (varying interstimulus intervals ranging between 70-330 ms between entrainers and 450-770 ms between entrainer 4 and the target).

**Figure 1:**
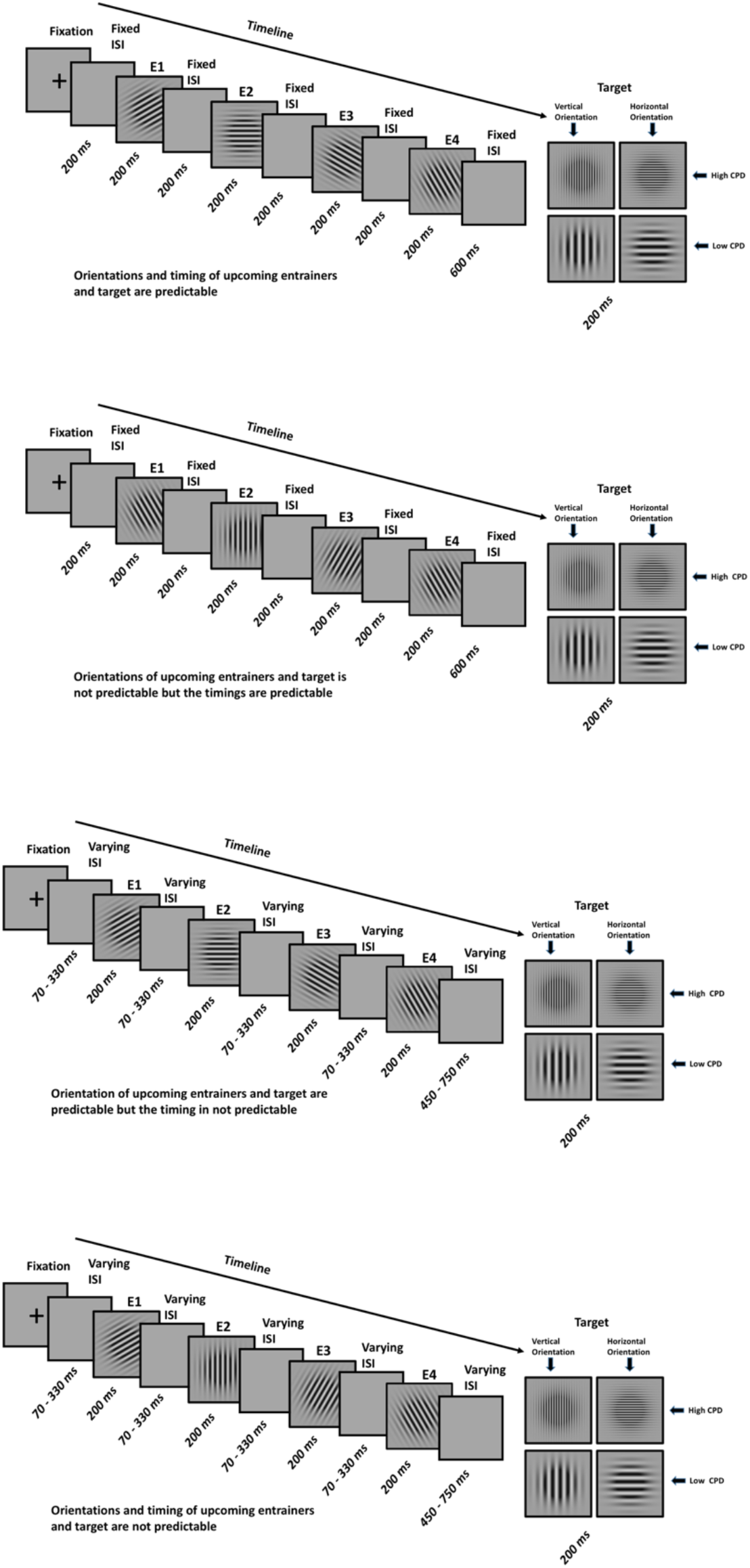
Experimental design. A) Orientations and timing of upcoming entrainers and target are predictable (*what*+*when* condition). B) Orientations of upcoming entrainers and target are not predictable but the timings are predictable (*when* condition). C) Orientation of upcoming entrainers and target are predictable but the timing is not predictable (*what* condition). D) Orientations and timing of upcoming entrainers and target are not predictable (*random* condition). Abbreviations: E1 – Entrainer 1, E2 – Entrainer 2, E3 – Entrainer 3, E4 – Entrainer 4, ISI – Inter Stimulus Interval

Depending on the timing and orientation of the entrainers and target, trials were divided into four conditions (Figure 1): (i) in the *what*+*when* condition, both the timing and the orientations of successive entrainers - and the final target Gabor - were predictable; (ii) in the *when* condition, timing was predictable but orientations were unpredictable. (iii) in the *what* condition, successive entrainers and target orientations were predictable but timing was unpredictable; (iv) in the *random* condition both orientations and timing were unpredictable.

A total of 160 trials were presented in each condition (80 horizontal and 80 vertical targets, randomly assigned 80 high and 80 low spatial frequencies) for a total of 640 trials per participant. 80 localizer trials for horizontal and vertical targets were also acquired while participants simply fixated the centre of the screen.

On each trial, participants had to indicate whether the target had a higher or a lower spatial frequency [CPD] than the preceding entrainers. Participants responded by pressing a button with their left or right hand, with the response hand counterbalanced across participants. A short optional break (participants pressed the button when they were ready to continue) was available after every 12 trials, and a longer mandatory break took place every 60 trials (the MEG researcher pressed a button from the operating console to pause and restart the presentation).

### Data acquisition and pre-processing

MEG data were acquired in a magnetically shielded room using the whole-scalp MEG system (Elekta-Neuromag, Helsinki, Finland) installed at the BCBL (http://www.bcbl.eu/bcbl-facilitiesresources/meg/). The system is equipped with 102 sensor triplets (each comprising a magnetometer and two orthogonal planar gradiometers) uniformly distributed around the head of the participant. Head position inside the helmet was continuously monitored using four Head Position Indicator (HPI) coils. The location of each coil relative to the anatomical fiducials (nasion, left and right preauricular points) was defined with a 3D digitizer (Fastrak Polhemus, Colchester, VA, USA). This procedure is critical for head movement compensation during the data recording session. Digitalization of the fiducials plus ∼300 additional points evenly distributed over the scalp of the participant were used during subsequent data analysis to spatially align the MEG sensor coordinates with T1 magnetic resonance brain images acquired on a 3T MRI scanner (Siemens Medical System, Erlangen, Germany). MEG recordings were acquired continuously with a bandpass filter of 0.01-330 Hz and a sampling rate of 1 kHz. Eye movements were monitored with two pairs of electrodes in a bipolar montage placed on the external canthus of each eye (horizontal electrooculogram (EOG)) and above and below the right eye (vertical EOG). Similarly, electrocardiogram (ECG) was recorded using two electrodes, placed on the right side of the participant’s abdomen and below the left clavicle.

Continuous MEG data were pre-processed off-line using the temporal Signal-Space-Separation (tSSS) method (Taulu & Simola, 2006) which suppresses external electromagnetic interference. MEG data were also corrected for head movements, and bad channel time courses were reconstructed in the framework of tSSS. Subsequent analyses were performed using Matlab R2014b (Mathworks, Natick, MA, USA).

### Behavioural data

Behavioural responses to the spatial frequency [CPD] task (on the target) were evaluated in terms of accuracy and Response Times (RTs) for all four conditions (*what+when, when, what, and random*). Trials with response times longer than 1500 ms were considered to be outliers and were removed from the analysis. The mean RT and standard deviation was computed for each experimental condition.

### Sensor level Event-Related Fields (ERFs)

MEG trials were corrected for jump and muscle artifacts using standard automated scripts based on the Fieldtrip toolbox (Oostenveld et al., 2011) implemented in MATLAB 2014B. Heartbeat and EOG artifacts were identified using Independent Component Analysis (ICA) and linearly subtracted from the MEG recordings. The ICA decomposition (30 components extracted per participant) was performed using the FastICA algorithm. ICA components maximally correlated with EOG and ECG recordings were automatically removed. On average, two components were removed per participant. The artifact-free data were bandpass filtered between 0.5 and 45 Hz. Trials were segmented time-locked to each of the entrainers (entrainers 1, 2, 3, and 4) and the target. The trial segments were grouped together for each entrainer and target, and then averaged to compute the ERFs. For each planar gradiometer pair, ERFs were quantified at every time point as the Euclidean norm of the two gradiometer signals. Baseline correction was also applied to the evoked data based on the 400 ms of data prior to the onset of the fixation cross presented at the beginning of each trial.

We applied an ANOVA to sensor-level data to explore the influence of our experimental factors on visual ERFs. First, we extracted ERF amplitudes in the set of five occipital sensors that had shown maximum response to the visual localizers. We then selected the time window classically associated with the initial visual evoked response (85-135 ms post stimulus). A three-way repeated measures ANOVA was computed in JASP (JASP Team, 2020) with these amplitude values as dependent variables and the following factors: entrainer (four levels; corresponding to entrainers 1, 2, 3, 4); *what* (two levels; predictable/unpredictable entrainer and target orientations); and *when* (two levels; predictable/unpredictable timing of entrainers and target).

Significant interactions (specifically, the triple interaction “entrainer * *what* * *when*”) were further investigated through theoretically relevant pairwise comparisons. Pairwise comparisons between conditions were performed using a cluster-based permutation test (Maris & Oostenveld, 2007). A randomization distribution of cluster statistics was constructed for each subject over time and sensors and used to evaluate whether conditions differed statistically over participants. In particular, t-values were computed for each sensor (combined gradiometers) and each time point during the 0-270 ms time window, and were clustered if they had t-values that exceeded a t-value corresponding to the 99.99^th^ percentile of Students t-distribution (*p* < 0.01 two tailed) and were both spatially and temporally adjacent. Cluster members were required to have at least two neighboring channels that also exceeded the threshold to be considered a cluster. The sum of the t-statistics in a sensor cluster was then used as the cluster-level statistic, which was then tested by permuting the condition labels 1000 times.

Four different comparisons were carried out. In the first comparison, we contrasted ERFs for the *when* and the *what*+*when* conditions. This comparison evaluated the effect of orientation predictability when the timing of the entrainers and target were predictable. In the second comparison, we compared ERFs for the *random* and *what* conditions. This comparison evaluated the effect of orientation predictability when timing was unpredictable. These two comparisons mainly focused on the main effect of orientation predictability (i.e., the *what* manipulation) revealed by expectation suppression.

We then compared the ERFs for the *what*+*when* and *what* conditions. Here we directly contrasted these two predictable orientation conditions to evaluate the effect of temporal predictability on stimulus predictability. The final comparison contrasted ERFs in the *when* and *random* conditions. This comparison was performed to analyse the effect of temporal predictability in the absence of orientation predictability.

### Source level Event-Related Fields (ERFs)

Source reconstruction mainly focused on the statistically more reliable effects observed at the sensor-level. In the present experimental scenario, we expected to find the strongest modulation of visual evoked responses for the last entrainer of each predictable series, when both *what* and *when* expectations would be highest.

MEG-MRI co-registration was performed using MRIlab (Elekta Neuromag Oy, version 1.7.25). Individual T1-weighted MRI images were segmented into scalp, skull, and brain components using the segmentation algorithms implemented in Freesurfer (Martinos Center of Biomedical Imaging, MQ; Dale et al., 1999). The source space was defined as a regular 3D grid with a 5 mm resolution and the lead fields were computed using a single-sphere model for 3 orthogonal source orientations. The lead field at each grid point was reduced to its first two principal components. Whole brain source activity was estimated using a linearly constrained minimum variance (LCMV) beamformer approach (Veen et al., 1997). Both planar gradiometers and magnetometers were used for inverse modelling. The covariance matrix used to derive LCMV beamformer weights was estimated from the pre- and post-stimulus data in the pre-stimulus (from 400 ms prior to fixation cross onset) to post-stimulus (400 ms after the presentation of the gabor) time range.

The LCMV beamformer focused on the (baseline corrected) evoked data in the time period 85–125 ms post-stimulus (when ERF peak amplitude across participants at the sensor level was largest). A non-linear transformation using the spatial-normalization algorithm (implemented in Statistical Parametric Mapping (SPM8), Friston et al., 1994) was employed to transform individual MRIs to the standard Montreal Neurological Institute (MNI) brain. Transformed maps were further averaged across participants. Freesurfer’s *tksurfer* tool was used to visualize the brain maps in MNI space. For each condition (at entrainer 4, E4), we obtained the source value and the MNI coordinates of local maxima (sets of contiguous voxels displaying higher source activation than all other neighbouring voxels; Bourguignon et al., 2018).

Source activity was compared between conditions (e.g., *when* vs. *what*+*when, random* vs. *what, what* vs. *what+when* and *when* vs. *random*) by extracting a peak value within a 5mm sphere around the common local maximum in the source space. We used t-tests to evaluate differences between conditions across participants.

### MVPA (Multivariate Pattern Analysis)

A MVPA approach (Pantazis et al., 2018; King et al., 2016; Cichy et al., 2014: Grootswagers et al., 2017) was used to evaluate the stimulus-specificity of the visual neural response across time in the experimental conditions. To validate our method we decoded both the feature of interest in the present experimental manipulation (i.e., the Gabor orientation) and a control feature of that same stimulus (i.e., its spatial frequency, or CPD).

Time-resolved within-subjects MVPA was performed to decode the features (i.e., the orientation and spatial frequency) of all the Gabors (i.e., E1, E2, E3, E4 and T) from the MEG data. For E1, E2, and E3, data were segmented from 50 ms prior to 250 ms after the onset of the entrainers. The time interval between E4 and the target was longer than the time interval between the rest of the entrainers. For this reason, for E4, the data was segmented from 50 ms prior to 600 ms after the onset of the entrainer. For the target, the data was segmented from −400 ms to 550 ms. The data were down sampled to 200 Hz prior to the classification procedure. Then the data were classified separately for both the orientation and spatial frequency of the Gabor using a linear support vector machine (SVM) classifier with L2 regularization and a box constraint of 1. The classifiers were implemented in MATLAB 2014B using the LibLinear package (Fan et al., 2008) and the Statistics and Machine Learning Toolbox (Mathworks, Inc.). We performed a binary classification of the orientation of each Gabor depending on the orientation and the spatial frequency of the subsequent target. In other words, the class labels (i.e., horizontal vs. vertical, higher vs. lower spatial frequency) were derived from the target orientation. For example, if the target orientation was horizontal, then all the preceding Gabor orientations in the corresponding condition were labelled as horizontal. A similar rationale was applied to classifying spatial frequency. Pseudo-trials were generated to improve SNR by averaging trials over bins of 10, without overlap (Dima and Singh, 2018). This pseudo-trial generation was repeated 100 times based on random ordering of the data to generate trials with a higher signal to noise ratio. The data were then randomly partitioned using 5-fold cross-validation. The classifier was trained on 4 folds and tested on the remaining fold; this process was repeated until each fold had been left out of the training once. The procedure of generating pseudo-trials, dividing the data into 5 folds, and training and testing classifiers at every time point was repeated 25 times; classification accuracies were then averaged over all these instances to yield more stable estimates. To improve classification, we also performed multivariate noise normalization (Guggenmos et al., 2018). The time-resolved error covariance between sensors was calculated based on the covariance matrix of the training set and used to normalize both the training and test sets in order to down-weight MEG channels with higher noise levels. Cluster corrected sign permutation tests (one-tailed) (Dima et al., 2018) were applied to the accuracy values obtained from the classifier with cluster-defining threshold *p* < 0.05, corrected significance level i.e., cluster-alpha *p* < 0.01.

## Results

### Behavioural results

On each trial, participants were asked to evaluate if the spatial frequency of the target (cycles per degree) was lower or higher than that of the entrainers. This stimulus dimension was not directly related to the experimental manipulation of timing (*when*) and Gabor orientation (*what*).

Table 1 presents the accuracy and reaction time (RTs) for all four conditions. We found no significant differences in behavioural accuracy. However, temporal predictability elicited faster RTs (*what+when* > *what* | *when* > *random*). We fit a Linear mixed model (*lmer*: R function) with participants and observations as *random* effects and *what* (orientation: predictable or not), *when* (timing: predictable or not) and their interaction as fixed effects. We observed an effect of *when* (t = –2.794, *p* < 0.05). Orientation predictability (*what*) did not elicit statistically significant effects (t = –1.557), probably due to that fact that, in order to perform the task, participants had to actively pay attention to the spatial frequency dimension of the target.

**Table 1:** Accuracy and Reaction Times (RTs) for each condition.

|  | *what+when* | *when* | *what* | *random* |
| --- | --- | --- | --- | --- |
| **Accuracy**<br>(Mean ± SD %) | 96.56 ± 3.49 | 95.89 ± 4.71 | 96.00 ± 3.80 | 96.56 ± 3.33 |
| **Reaction Times**<br>(Mean ± SD ms ) | 727.5 ± 204 | 737.4 ± 203 | 741.4 ± 204 | 752 ± 207 ms |

**Table 2:**
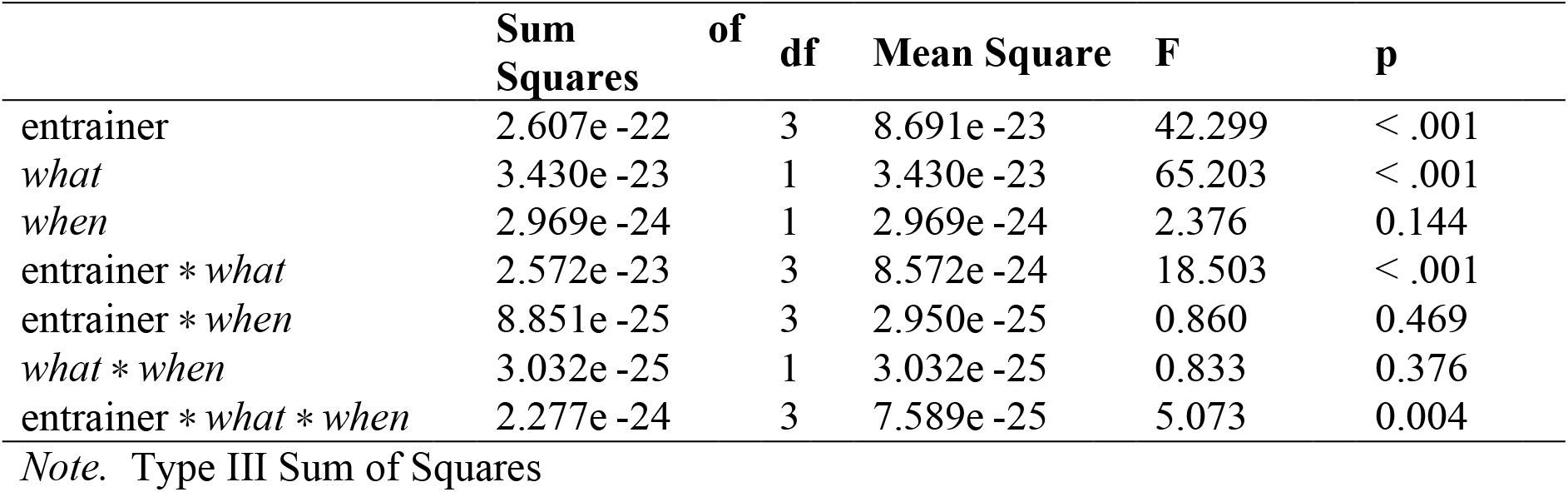
Repeated measure ANOVA with the factors entrainer (four levels, one for each entrainer), what (two levels: orientation predictable or not) and when (two levels: timing predictable or not).

**Table 3:** Statistically significant classification accuracy for orientation angle and spatial frequencies (CPD) of the target Gabors across entrainers and target for the orientation predictable conditions. Time intervals indicate the windows in which accuracy was statistically above chance. n.s.: not significant.

| Feature | Condition | E1 | E2 | E3 | E4 | Target |
| --- | --- | --- | --- | --- | --- | --- |
| **Orientation angle** | *what+when* | 135 - 150 ms<br>(56.82 %) | -20 - -10 ms<br>(57.41 %) | 100 - 125 ms<br>(63.07 %) | 95 - 215 ms<br>(70.46 %) | 95 - 450 ms<br>(78.95 %) |
|  | *what* | 140 - 165 ms<br>(56.88 %) | n.s. | 125 - 160 ms<br>(60.20 %) | 100 - 170 ms<br>(68.32 %) | 95 - 275 ms<br>(75.29 %) |
| **Cycles per degree (CPD)** | *what+when* | n.s. | n.s. | n.s. | n.s. | 80 - 550 ms<br>(97.73 %) |
|  | *what* | n.s. | n.s. | n.s. | n.s. | 85 - 550 ms<br>(97.19 %) |

### Sensor-level MEG results

Next, we report the analysis of evoked responses to the four entrainers, where visual predictions were incrementally built up. Note that there was no explicit task related to orientation. We first analysed the amplitude of the initial visual evoked response for 5 occipital sensors (the ones showing maximum response to the localizers) to determine how the two-by-two experimental design modulated visual responses across entrainers. In the three-way ANOVA (details in Table 1), we observed a significant main effect of entrainer (*p* < 0.001) on the peak amplitudes of ERFs. Interestingly, this factor interacted with the factor *what* (*p* < 0.001), suggesting that Gabor orientation predictability affected visual-evoked responses differently across entrainers. We should point out that a main effect of *what* (*p* < 0.001) supported the observation that orientation predictability influenced visual processing. Importantly, the interaction between the three factors, i.e., entrainers, *what* and *when*, was significant (*p* = 0.004). This triple interaction underlines the fact that timing uncertainty influenced the development of visual predictions across the sequence of four entrainers.

Figure 2 shows the sensor-level ERFs time-locked to the onset of each entrainer (E1, E2, E3, and E4) and target (T) for the *when* and the *what+when* conditions. Here, we can see the influence of orientation predictability (expectation suppression effect) when timing was also predictable. The amplitude of the ERFs was significantly higher (*p* < 0.01, cluster based permutation test) in the *when* than the *what+when* condition for E2, E3, and E4, but not for E1 and T. The amplitude enhancement for the *when* compared to the *what+when* condition emerged in the (expected) 95–105 ms, 96–110 ms, and 97–121 ms time intervals for E2, E3, and E4, respectively. These clusters were located in occipital sensors for all four entrainers.

**Figure 2:**
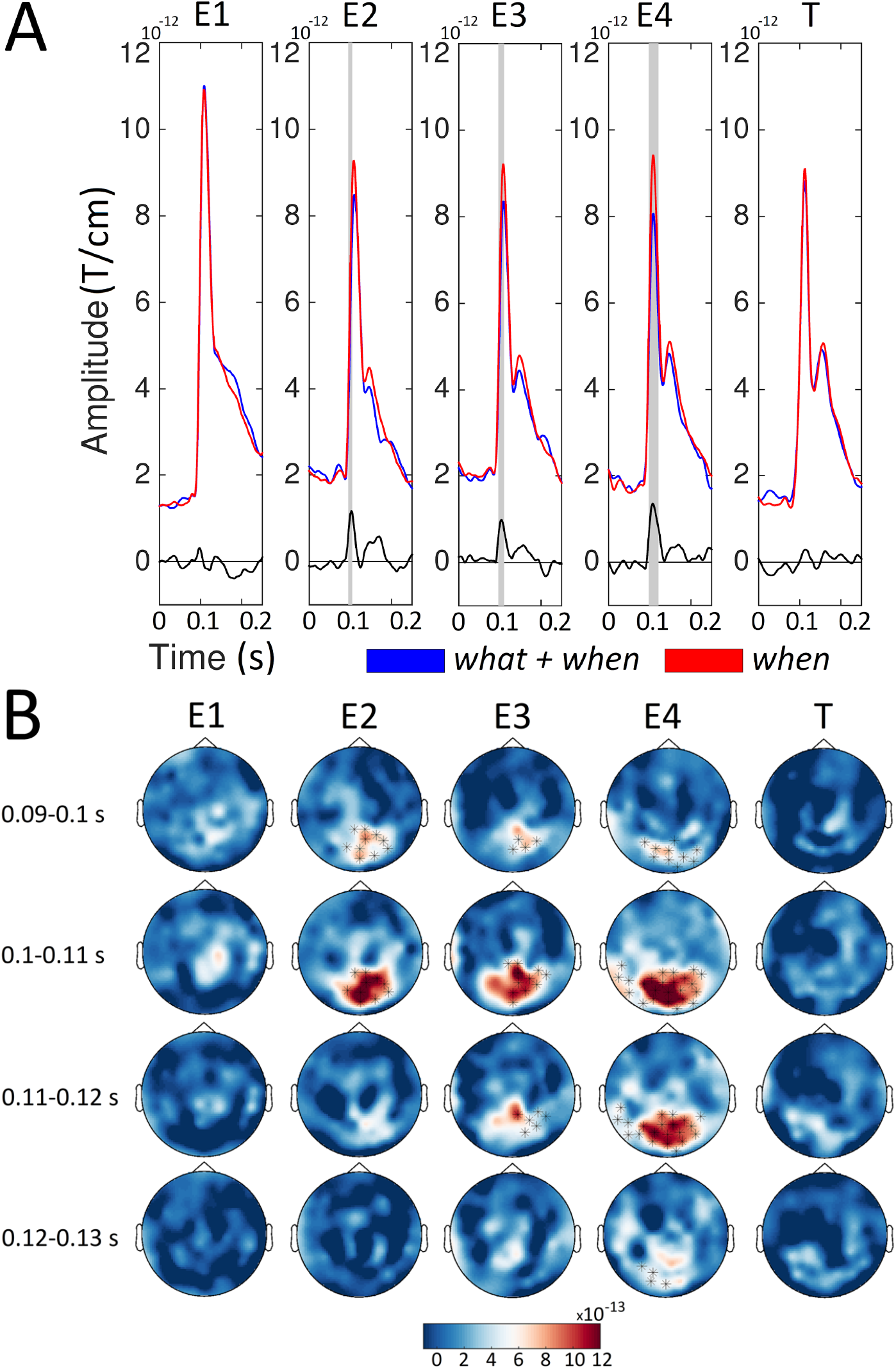
Sensor level ERFs for the *when* and *what*+*when* conditions. A) For each condition (red, *when*; blue, *what*+*when*) and stimulus (Entrainer 1 [E1], E2, E3, E4 and Target [T]), we show the average of the event related fields (ERFs) in representative channels located above occipital regions (MEG02042/3, MEG2032/3, MEG2342/3, MEG2122/3, and MEG1922/3). Below we also report the ERF difference between the *when* and *what*+*when* (black line) conditions. Grey boxes represent time points where the amplitude of the ERFs was higher (*p* < 0.01, cluster-based permutation test) for the *when* than the *what*+*when* condition. B) Sensor maps of the ERF difference between the *when* and *what*+*when* conditions in temporal windows ([0.090 – 0.100], [0.100 – 0.110], [0.110 – 0.120] and [0.120 – 0.130] s) around the amplitude peak value. Sensors showing significant differences (*p* < 0.01, cluster-based permutation test) are highlighted.

Figure 3 shows the sensor-level ERFs for the *random* versus the *what* conditions. This comparison highlights the expectation suppression effect when timing was not predictable. The amplitude of the ERFs was significantly higher (*p* < 0.01) for the *random* compared to the *what* condition for E2, E3, and E4 but not for E1 and T. The amplitude enhancement for the *random* compared to the *what* condition emerged within the 95–119 ms, 94–123 ms, and 96–127 ms time intervals for E2, E3, and E4, respectively. These clusters were also clearly distinguishable in occipital sensors for all the entrainers.

**Figure 3:**
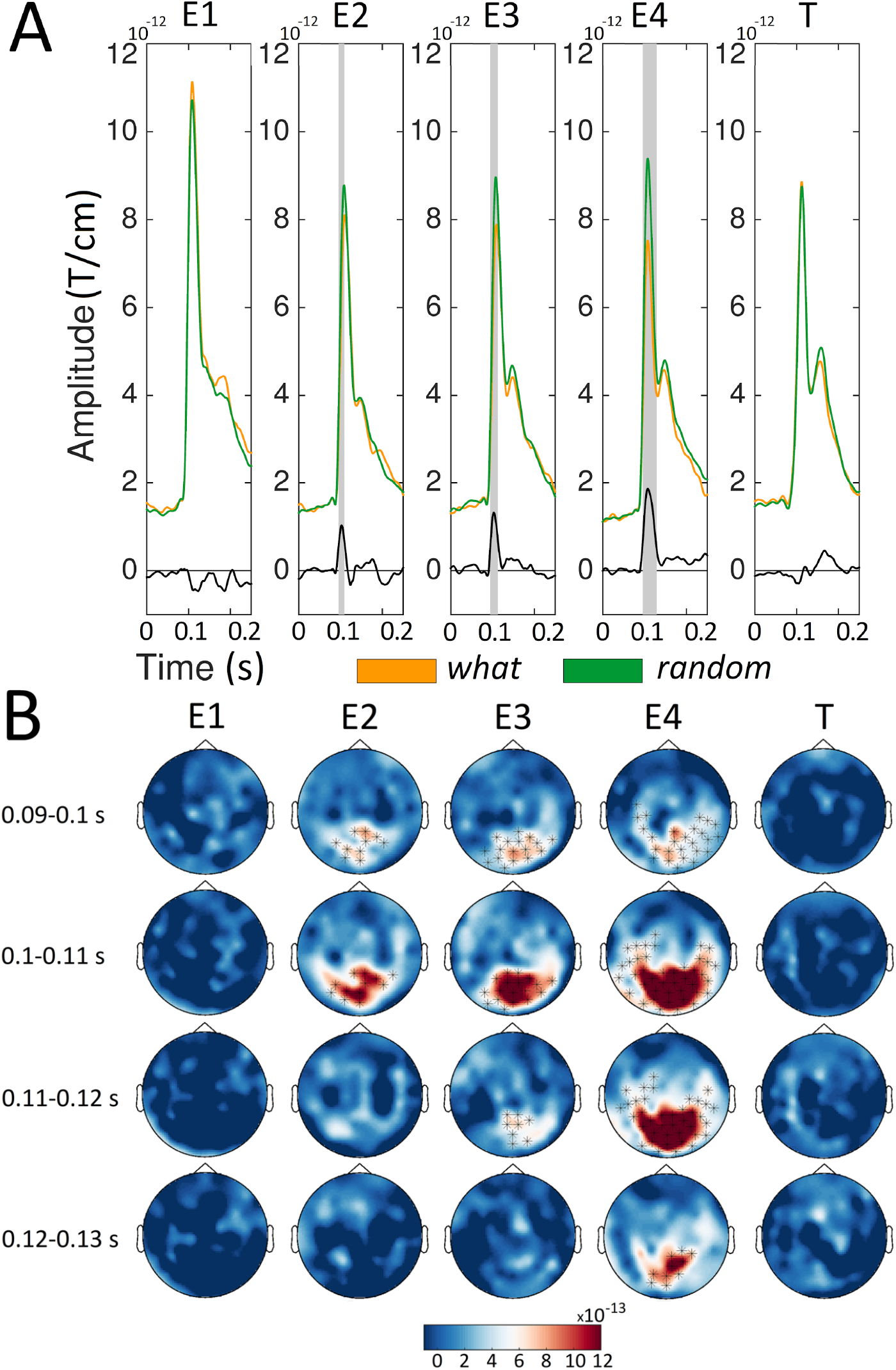
Sensor level ERFs for the *what* and *random* conditions. For each condition (orange, *what*; green, *random*) and stimulus (Entrainer 1 [E1], E2, E3, E4 and Target [T]), we showed the average of the Event Related Fields (ERFs) Grey boxes represent time points where the amplitude of the ERFs was higher (*p* < 0.01, cluster-based permutation test) for the *random* than the *what* condition. B) Sensor maps of ERF differences between the *random* and *what* conditions in temporal windows ([0.090 – 0.100], [0.100 – 0.110], [0.110 – 0.120] and [0.120 – 0.130] s) around the amplitude peak value. Sensors showing significant differences (*p* < 0.01, cluster-based permutation test) are highlighted.

Since both comparisons involving expectation suppression were significant from entrainer 2 onward, we compared the two orientation predictable conditions with (*what+when*) and without (*what*) temporal predictability. This comparison highlights how temporal predictability affects visual predictive processing. Figure 4 shows that initial early evoked activity at E1 (0–75 ms, preceding the peak reflecting the visual evoked response) is similar for both conditions. As we move across entrainers, these early differences increase and reach statistical significance (*p* < 0.001), but this effect vanishes at the target. This differential pre-stimulus activity demonstrates that results for the two orientation predictable conditions depend on temporal predictability. Here, it is worth noting that we used the same baseline time period (the 400 ms before the fixation cross at the beginning of the whole trial) to test the effects of all four entrainers (E1 to E4) and the target (T). Since our focus in this analysis was on early evoked responses to the visual stimulus, which showed robust expectation suppression effects across all predictable entrainers (see Figures 2 and 3), we selected the time window from 75 to 135 ms, corresponding to the initial ERF peak reflecting early visual processing, for statistical comparison. Across the four entrainers an effect emerged only at E4, where the amplitude suppression was larger for the *what* than for the *what+when* condition in a cluster spanning the 106–124 ms time interval. This cluster was located in occipital sensors (Figure 4).

**Figure 4:**
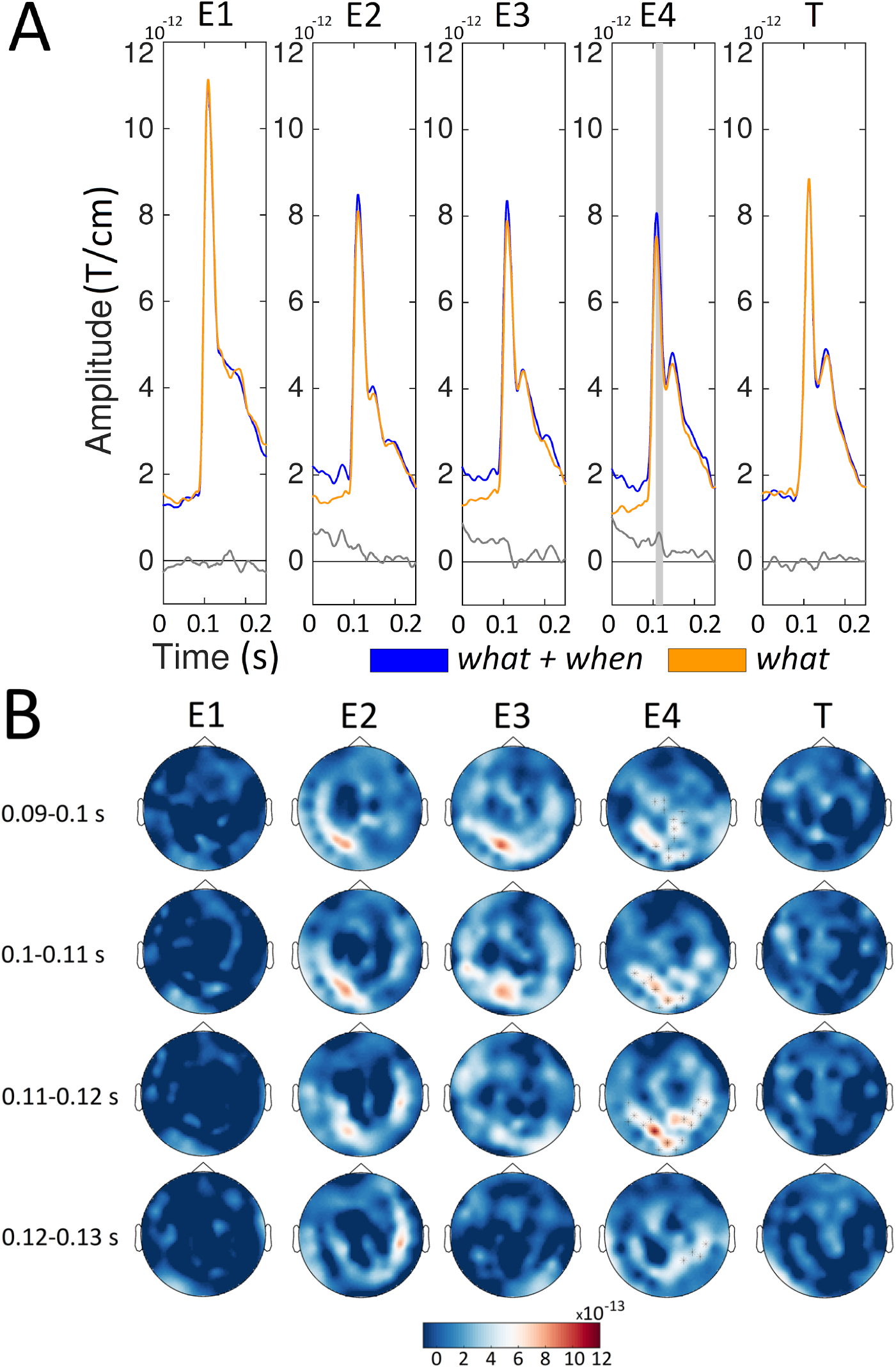
Sensor level ERFs for the *what*+*when* and *what* conditions. A) For each condition (orange, *what*; blue, *what*+*when*) and stimulus (Entrainer 1 [E1], E2, E3, E4 and Target [T]). Grey boxes represent time points where the amplitude of the ERFs (in the [0.075 – 0.135] s window) was higher (*p* < 0.05, cluster-based permutation test) for the *what*+*when* than the *what* condition. B) Sensor maps of the ERF difference between the *what*+*when* compared to the *what* condition in temporal windows ([0.090 – 0.100], [0.100 – 0.110], [0.110 – 0.120] and [0.120 – 0.130] s) around the amplitude peak value. Sensors showing significant differences (*p* < 0.01, cluster-based permutation test) are highlighted.

The differences that emerged between the two orientation predictable conditions (Figure 4) across the entrainment sequence could be due to carry-over effects from an earlier difference in baseline activity. To evaluate this hypothesis we also compared the two orientation unpredictable conditions (*when* and *random*). Here, we expected similar differential baseline activity in the 0–75 ms time interval, but no difference at the peak of the visual response for any entrainer. Figure 5 shows the sensor level comparison of the ERFs for the *when* and *random* conditions. Differences in the initial activity time-locked to the Gabor patch are evident at E2, E3, and E4 within the 0–75 ms time range (*p* < 0.001, compare Figures 4 and 5). This difference is not evident at E1 or T. Since evoked responses to all the entrainers (E1, E2, E3, and E4) and T were baseline corrected using the same activity period before the fixation cross at the beginning of the trial, this effect could reflect temporal predictability affecting ongoing brain activity before the initial visual response to each Gabor entrainer. Importantly, we focused on the effect of temporal predictability on the peak visual response. In the selected time window 75–135 ms we did not observe any statistically reliable effect at any entrainer. This null effect is in line with the triple interaction observed in the initial overall ANOVA (Table 1), supporting the idea that temporal predictability affects visual predictive processing.

**Figure 5:**
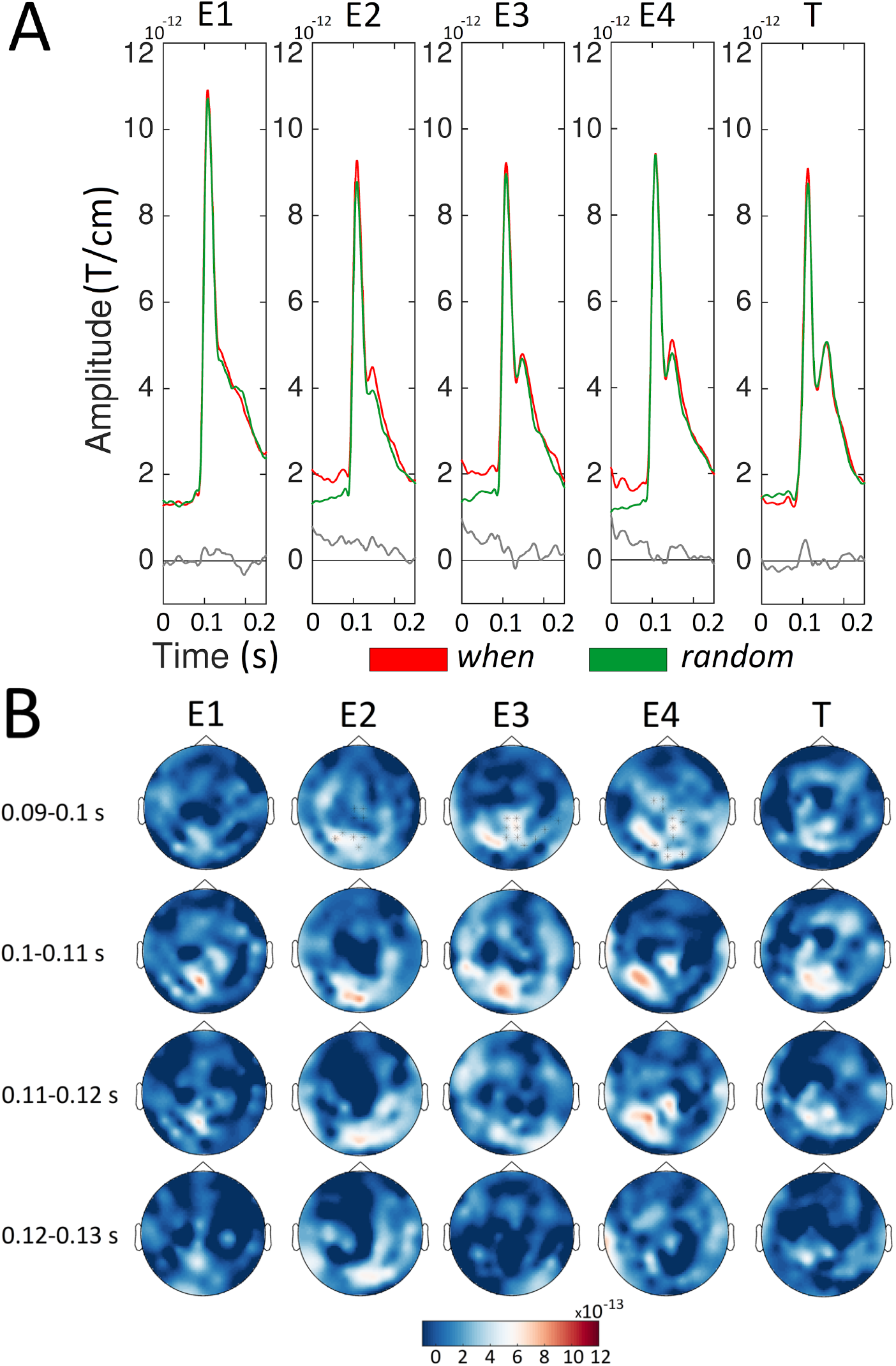
Sensor level ERFs for the *when* and *random* conditions. A) For each condition (red, *when*; green, *random*) and stimulus (Entrainer 1 [E1], E2, E3, E4 and Target [T]). B) Sensor maps of the ERF differences between the *when* and the *random* conditions in temporal windows ([0.090 – 0.100], [0.100 – 0.110], [0.110 – 0.120] and [0.120 – 0.130] s) around the amplitude peak value. Sensors showing significant differences (*p* < 0.01, cluster-based permutation test) are highlighted.

### Source level ERFs

We next identified the brain regions underlying the relevant effects observed at the sensor level. Source activity was estimated around the peak amplitude of the sensor-level ERFs in the 85–125 ms interval. Whole-brain maps of source activity were created for each condition (*what*+*when, when, what* and *random*) and entrainer 4 (E4), i.e., the stimulus where the difference between *what*+*when* and *what* was statistically significant (Figure 6).

**Figure 6:**
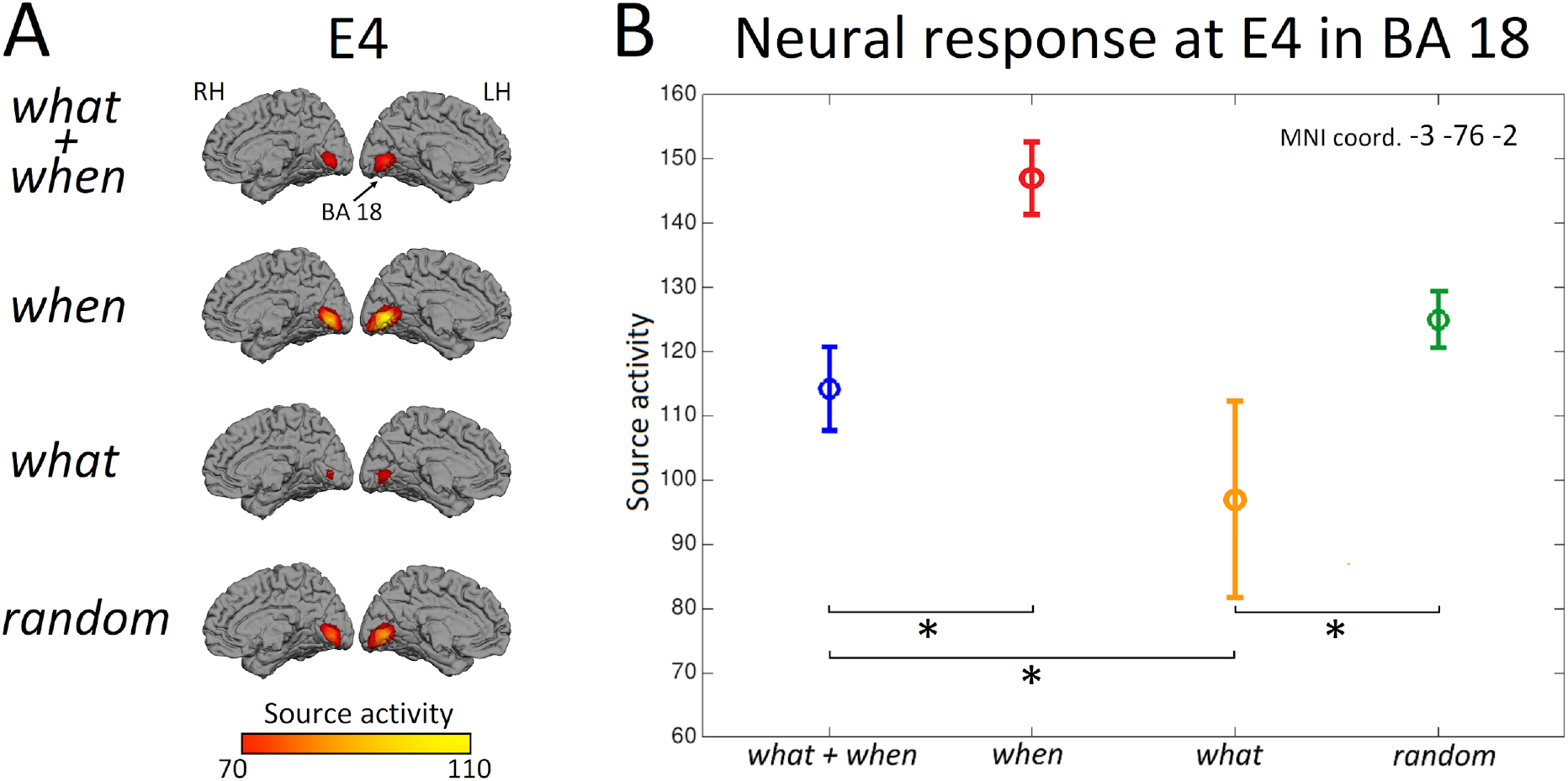
A) Brain maps representing source activity for each condition (*what*+*when, when, what*, and *random*) at Entrainer 4 (E4). We included a view of the medial surface and the occipital lobe of the left (LH) and the right (RH) hemispheres. B) The mean source activity in BA18 (Brodmann Area 18: xyz MNI coordinate : –3, –76, –2) in the four conditions at E4. Asterisks indicate significant differences across conditions.

Source activity was localized in bilateral occipital regions for all conditions and compared to baseline at the group level. The first local maxima emerged in visual association areas (Brodmann Area 18: BA 18, average coordinate [–3, – 76, –2]) of the left occipital cortex in all conditions.

For this local maximum we evaluate the amplitude of source activity across conditions, following the same rationale described for the sensor-level analyses. Figure 6A shows the brain maps representing maximum peak activity in the source space. Figure 6B shows source amplitudes with the corresponding standard error. Source amplitude was significantly higher for the *when* than the *what*+*when* condition (*t* = 4.16, *p* < 0.05) and for the *random* compared to the *what* condition (*t* = 5.20, *p* < 0.05) at E4 . Crucially, these values were higher for the *what+when* than the *what* condition (*t* = 2.38, *p* < 0.05), while no difference emerged between the *when* and *random* conditions. Overall, the present results confirm the effects observed at the sensor-level, providing a candidate location for the generation of the expectation suppression effects reported at the sensor level.

### MVPA results

The ERF analyses showed that the expectation suppression effect grew incrementally larger across the four entrainers, demonstrating that orientation predictability reduced visual processing costs, possibly due to increased reliance on internally generated expectations. To further corroborate the hypothesis that the visual system developed expectations for successive Gabor orientations during the entrainer sequence, we performed the following analyses. We first checked whether the orientation of perceived Gabors could be decoded by applying a temporal decoding approach to each entrainer (as a control we performed the same analysis to decode the spatial frequencies of these Gabors). MVPA showed that only those conditions with predictable orientations (*what+when* and *what*) revealed above-chance and statistically significant decoding accuracy values compared to the conditions where orientation was not predictable (see Supplementary Figures 1 and 2 for the comparison *what+when* vs. *when* and *what* vs. *random*, respectively). Figure 7 shows the decoding accuracy of predictable orientations in conditions with (*what+when*) and without (*what*) temporal predictability. Here, we see how target orientation becomes increasingly decodable across entrainers (especially at E3 and E4; t-test between peak decoding accuracy at E1 and E4 across participants and conditions, *p* < 0.01) and is strongest at the target. By contrast, decoding values for spatial frequencies (high vs. low CPD) were significant only at the target. This was expected since the CPD of the target could be predicted from the entrainers, which had intermediate spatial frequencies; this provided a good baseline for target orientation decoding effects.

**Figure 7:**
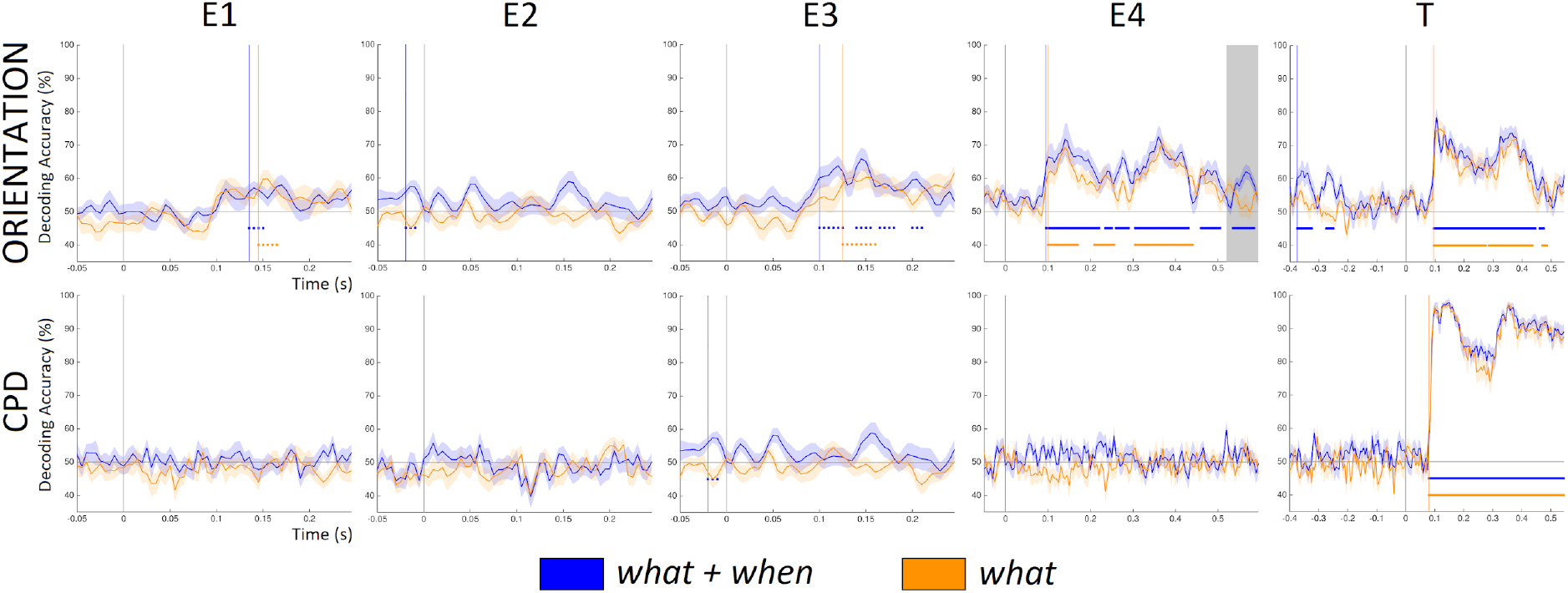
Time-resolved decoding accuracy for the *what*+*when* condition (blue line) and *what* condition (orange line) time-locked to Entrainer 1 (E1), E2, E3, E4 and Target (T). The coloured dots under the curves indicate the statistical significance of decoding accuracy across time. The gray box at E4 in Orientation shows the statistical significant differences (*p* < 0.05) between *what+when* and *what*.

Notably, decoding accuracy (at E3 and E4) was reliable within the time period of the initial evoked response (75–200 ms) for the two orientation predictable conditions whether timing was predictable or not. No statistical difference between the two predictable conditions emerged in this time interval. This indicates that the neural representation of the visual stimulus was preserved independently of the amplitude of the ERF responses. This analysis also that decoding accuracy was higher for Entrainer 4 in a later time interval (525– 595 ms, *p* < 0.05) for the *what*+*when* than the *what* condition. This indicates that visual representations were actively maintained for a longer time period when timing was isochronous, while the effect lasted for less time when timing was jittered.

It could be argued that the increase in decoding accuracy for the orientation of the target observed across entrainers could be due to the fact that entrainer orientations gradually approached a horizontal/vertical orientation, making them representationally more similar to the target. . We argue that this is not the case based on two facts. For one. if this had been the case, we should have found large decoding accuracy even at the first entrainer. The orientation of the first (compared to following) entrainer was closer to the horizontal vs. vertical orientation contrast, while being orthogonal to the target (if E1 was horizontal, the target would be vertical). The classifier should also have picked up this dissociation at the first entrainer (since the classifier was blind to the actual orientation of the stimuli in any two classes and was simply trained to evaluate whether two classes of data were different). This was not the case (with weak and unstable accuracy for E1), indicating that the classifier was instead picking up increasing “orientation expectation” across the sequence of entrainers. Second, if the classifier had detected overall visual similarity (not specific to orientation) between entrainers and the target, we would have expected a similar trend of increasing accuracy to also emerge in decoding spatial frequencies (CPD). However, at E4 decoding accuracies for spatial frequency consistently showed chance level accuracy.

## Discussion

In the present study we report robust expectation suppression effects for predictive processing of visual Gabors. Across a series of four entrainers, we observed incrementally larger suppression of visual evoked responses when Gabor orientations were predictable, accompanied by incrementally improved decodability for predictable Gabor orientations. Importantly, these effects were modulated by temporal predictability: expectation suppression of evoked responses was larger when the timing of the entrainers was jittered, while decodability of visual responses was less sustained for jittered timings. These findings indicate that the neurocognitive system invested less resources in visual analysis in temporally uncertain scenarios, possibly due to higher reliance on internal predictions.

### Expectation suppression effects

The goal of our study was to evaluate how visual expectations are modulated by temporal predictability. We mainly focused on the expectation suppression effect (Walsh et al., 2020), and in contrast to previous studies (Auksztulewicz et al., 2018; Utzerath et al., 2017), (i) we did not use mismatched stimuli, and (ii) we made sure that participants were not aware of the experimental manipulations in the study. Given the much debated interaction between attention and predictive processing (Kok et al., 2012), we developed an experimental design which aimed to control for strategic effects related to the processing of the Gabor orientation (the task required that participants instead focus on spatial frequencies, which were the same across conditions). While the orientation manipulation was noticeable, it is important to underscore that our participants did not report having observed any temporal jitter of the visual stimuli in the temporally unpredictable conditions.

There are several studies in which reduced neural responses for predictable stimuli have been found during passive viewing (Alink et al., 2010), as well as when stimuli are fully task irrelevant (Den Ouden et al., 2009). This supports the contention that expectation suppression does not vary whether or not participants engage with the task. However, some authors have reported no effects from expectation suppression of sensory activity when stimuli were unattended (Larsson and Smith, 2012). These findings suggest that contextually predictable stimuli may not result in any suppression of early visual neural responses (John-Saaltink et al., 2015). In the present study, we found expectation suppression effects when the orientation of the entrainers was predictable and showed that these effect increased incrementally across the entrainer sequence. We interpret this effect as demonstrating that the visual system develops increasingly strong expectations for the specific orientation of upcoming Gabors across sequences: the stronger the expectation for a Gabor orientation, the larger the suppression of the visual response. This effect was significant in the evoked responses of the second, third and fourth entrainers (Figures 2 and 3) and was present in conditions both with and without temporal predictability.

This visual evoked response possibly originated in visual area 2 (V2), the area which showed reduced activity for predictable stimuli compared to unpredictable stimuli (Figure 6). The source location of the present effect could reflect some sort of top-down activity generated in an extrastriate region projecting to primary visual cortex (V1). This possibility should, however, be further validated (possibly by employing direct brain recordings in non-human primates) with additional connectivity analyses to investigate the bidirectional interaction between V1 and V2 and determine if the flow of information in the top-down direction is enhanced for content predictable conditions.

It is worth noting that this incremental effect was not mirrored in the behavioural responses, which probably reflect later decision processes. In addition, expectation suppression effects evident during the entrainer sequence vanished at the presentation of the target Gabor (where participants had to perform the task). At the target, it is possible that neural resources were largely invested in processing the task-relevant spatial frequency difference between the target and the preceding entrainer Gabors. This task likely interacted with and washed out the on-going neural expectation effects, which were observed for entrainers. It would be interesting to evaluate how the different features of the target Gabor (orientation and spatial frequency) were processed in future studies. This could be achieved by using a delayed cueing task in which participants receive a random post-target cue after target presentation indicating whether they should perform an orientation or spatial frequency discrimination task. Another option would be to avoid the use of any task, i.e., employ a passive viewing paradigm, to determine if the expectation suppression effect is nevertheless preserved.

### The interaction between stimulus features and temporal predictability

We observed a repetition suppression effect in evoked responses within an early time interval, i.e., at 85–125 ms. The analysis of the amplitude of this initial evoked response showed that there was a significant interaction between the *what* and *when* dimensions of visual stimuli. This interaction was mainly driven by the increased suppression of neural responses to temporally jittered vs. isochronous predictable stimuli. It seems that the visual system generates a reduced response to an incoming stimulus whose onset is unpredictable compared to a stimulus whose timing is certain. It could be argued that the visual system is not “capable of preparing” for a temporally unpredictable stimulus. This can be supported by the fact that the effects observed in the evoked peaks at around 100 ms are preceded by a large difference between temporally predictable and unpredictable conditions independently from stimulus content predictability (Figures 4 and 5).

However, at around 100 ms, when the initial visual evoked response is peaking, a difference emerges between the two content predictable conditions (*what+when* vs. *what*, Figure 4) but not between the two content unpredictable conditions (*when* vs. *random*, Figure 5; see also Figure 6). If the reduced response found for temporally unpredictable stimuli in the *what* condition (compared to *what*+*when*) had only been driven by the inability of the visual system to prepare for the timing of a visual stimulus (regardless of its visual properties), one would have expected a similarly larger visual response in the *random* condition (compared to *when*) as well. One potential explanation for the larger suppression of the early evoked response in the temporally unpredictable *what* condition is that the visual system relies more on internal predictions and less on external evidence, when temporal uncertainty is higher (if predictions have been developed). This evidence would support theoretical claims suggesting that predictive mechanisms are essential for reducing uncertainty about the external environment (Clark, 2013).

### Stimulus specific neural activity

Expectation suppression effects provided evidence for reduced visual processing for expected stimuli, but did not provide evidence for expectations of specific visual representations. We thus used multivariate pattern analysis to evaluate whether expectations regarding Gabor orientation increased across entrainers. Our results show that decoding accuracy for Gabor orientations increased across entrainers when successive entrainer and target orientations were predictable (Figure 7). This indicates that stimulus predictability is a crucial factor in enhancing the accuracy of orientation decoding during the presentation of entrainers. In fact, when the stimulus is not predictable, decoding accuracy remains at chance level (Supplementary Figures 1 and 2).

Temporal predictability did not affect the decodability of the predicted visual stimulus in the earlier time interval, when the early visual evoked response emerged. This indicates that the representation of the Gabor orientation was stable and preserved independently of the amplitude of the related evoked response. On the other hand, temporal predictability differently affected orientation decoding in a later time interval (525–595 ms), showing that the orientation representation of the fourth entrainer was maintained active for a longer period of time if the timing of the stimulus was predictable. In other words, this suggests that the visual system invests more resources and prolongs processing of stimulus features when the temporal onset of the visual stimulus is predictable. This difference mirrors the evoked effects, where we observed stronger visual responses to temporally predictable than temporally unpredictable conditions. The decoding results thus reinforce our hypothesis that stimulus-specific neural processes are recruited to a larger extent for processing expected/temporally predictable visual stimuli, compared to expected/temporally uncertain visual stimuli.

A side note regarding the decoding of spatial frequency: this feature was constant across entrainers and conditions thus leading to chance level decoding for all four entrainers. At the target, however, spatial frequency showed very high decoding accuracy (around 97–98 %), even higher than orientation decoding, within the time interval related to the initial evoked response (∼100 ms). Spatial frequency effects were also evident slightly earlier and lasted longer than the orientation effects. Since our task focused on the difference in spatial frequency between the entrainers and target, the neural system likely maintained information regarding the spatial frequency of the target active for a longer interval that information regarding the orientation of the target, thus obscuring or interfering with any on-going expectation suppression effects due to Gabor orientation.

### Conclusions

In the present study we investigated the effect of temporal predictability on visual predictive processing. Our results show that temporal predictability modulates processing of expected visual features. We found increased suppression of visual evoked responses for temporally unpredictable relative to temporally predictable visual stimuli. This may demonstrate that the brain increases reliance on more abstract internal and representations when timing is uncertain.

## Acknowledgements

This research was supported by the Basque Government through the BERC 2018-2021 program and by the Spanish State Research Agency through BCBL’s Severo Ochoa excellence accreditation SEV-2015-0490 and through the project BES-2016-077560 funded by the Spanish Ministry of Economy and Competitiveness (MINECO). SN acknowledges support from an EMBO short term fellowship. NM was supported by the Spanish Ministry of Science, Innovation and University (grants PSI2015-65694-P, RTI2018-096311-B-I00), the Agencia Estatal de Investigación (AEI), the Fondo Europeo de Desarrollo Regional (FEDER) and by the Basque government (grant PI_2016_1_0014). RMC was supported by the German Research Council (DFG) (CI241/1-1, CI241/3-1) and the European Research Council (ERC-StG-2018-803370). The authors thank Asier Zarraza, Ning Mei, David Soto, and Lucia Amoruso for their valuable comments on the initial draft of this publication. SN thanks the BCBL lab research staff for their valuable support.

## Declaration

The **author**(s) **declare**(s) that there is **no conflict of interest** regarding the publication of this article.

## Supplementary figures

**Supplementary Figure 1:**
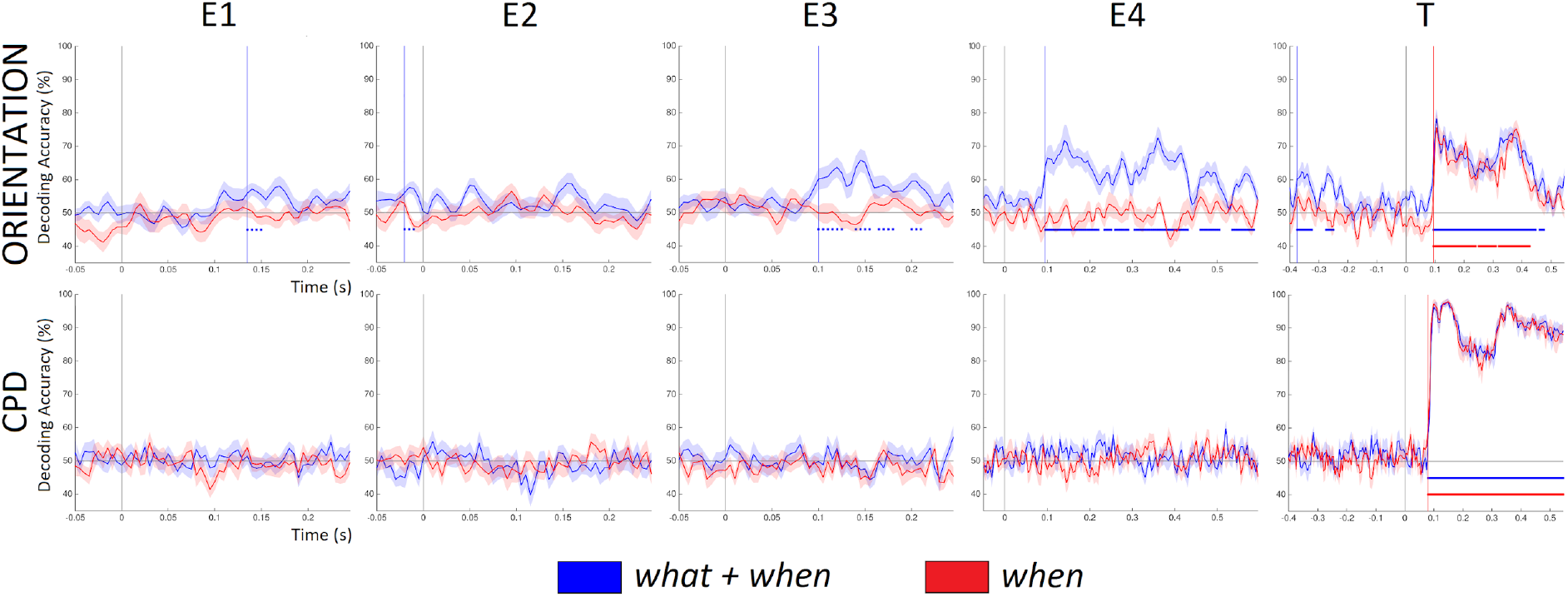
Time resolved decoding of *what*+*when* and *what* trials time-locked to Entrainer 1(E1), Entrainer 2 (E2), Entrainer 3 (E3), Entrainer 4 (E4) and Target (T). The coloured dots represent the statistical significance of accuracy.

**Supplementary Figure 2:**
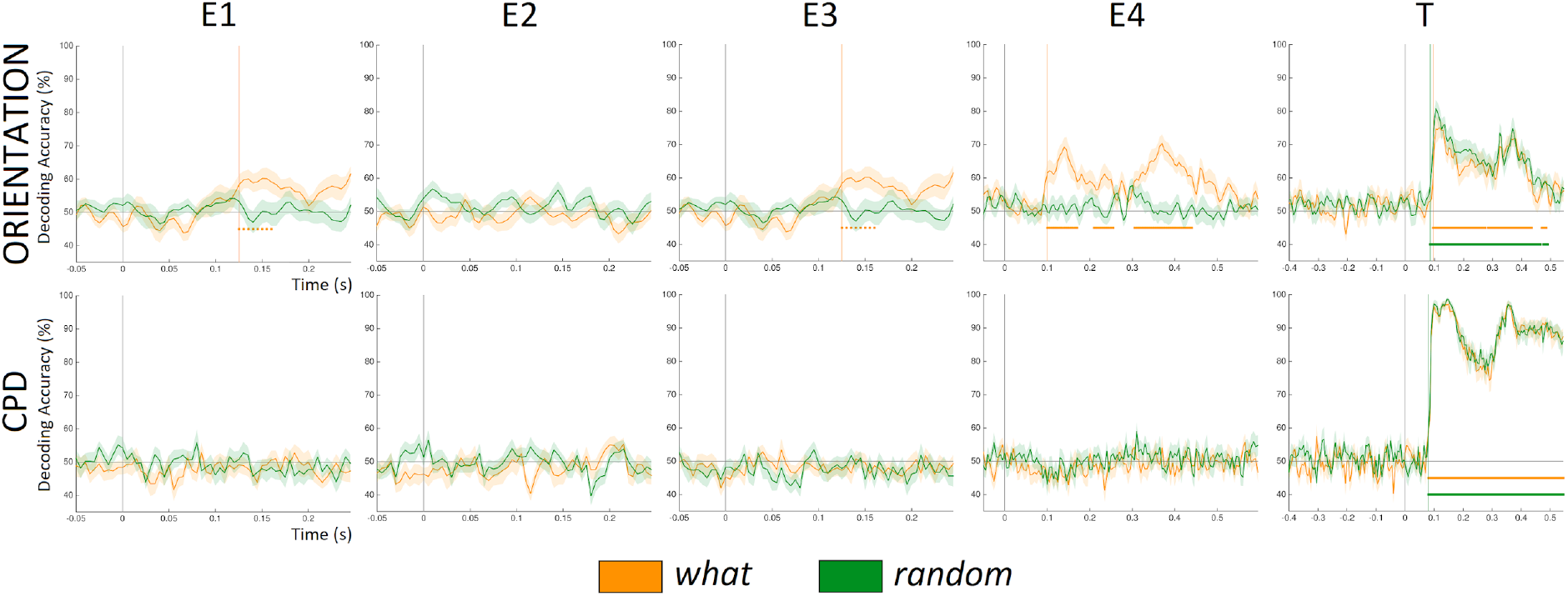
Time resolved decoding of *what* only and *random* trials time-locked to Entrainer 1(E1), Entrainer 2 (E2), Entrainer 3 (E3), Entrainer 4 (E4) and Target (T). The coloured dots represent the statistical significance of accuracy.

**Supplementary Table 1.**

| **Feature** | **Condition** | **E1** | **E2** | **E3** | **E4** | **Target** |
| --- | --- | --- | --- | --- | --- | --- |
| **Orientation angle** | *what+when* | 135 - 150 ms<br>(56.82 %) | -20 - -10 ms<br>(57.41 %) | 100 - 125 ms<br>(63.07 %) | 95 - 215 ms<br>(70.46 %) | 95 - 450 ms<br>(78.95 %) |
|  | *when* | ** | ** | ** | ** | 95 - 240 ms<br>(76.84 %) |
|  | *what* | 140 - 165 ms<br>(56.88 %) | ** | 125 - 160 ms<br>(60.20 %) | 100 - 170 ms<br>(68.32 %) | 95 - 275 ms<br>(75.29 %) |
|  | *random* | ** | ** | ** | ** | 85 - 465 ms<br>(75.39 %) |
| **Cycles per degree (CPD)** | *what+when* | ** | ** | ** | ** | 80 - 550 ms<br>(97.73 %) |
|  | *when* | ** | ** | ** | ** | 80 - 545 ms<br>(97.92 %) |
|  | *what* | ** | ** | ** | ** | 85 - 550 ms<br>(97.19 %) |
|  | *random* | ** | ** | ** | ** | 80 - 545 ms<br>(98.65 %) |

\*\* : No Significant cluster found.

## References

Alink, A., Schwiedrzik, C.M., Kohler, A., Singer, W., Muckli, L., 2010. Stimulus predictability reduces responses in primary visual cortex. J. Neurosci. 30, 2960–2966. https://doi.org/10.1523/JNEUROSCI.3730-10.2010

Auksztulewicz, R., Schwiedrzik, C.M., Thesen, T., Doyle, W., Devinsky, O., Nobre, A.C., Schroeder, C.E., Friston, K.J., Melloni, L., 2018. Not all predictions are equal: “what” and “when” predictions modulate activity in auditory cortex through different mechanisms. J. Neurosci. 38, 8680–8693. https://doi.org/10.1523/JNEUROSCI.0369-18.2018

Blom, T., Feuerriegel, D., Johnson, P., Bode, S., Hogendoorn, H., 2020. Predictions drive neural representations of visual events ahead of incoming sensory information. https://doi.org/10.1073/pnas.1917777117

Bourguignon, M., Molinaro, N., Wens, V., 2018. Contrasting functional imaging parametric maps: The mislocation problem and alternative solutions. Neuroimage 169, 200–211. https://doi.org/10.1016/j.neuroimage.2017.12.033

Carlson, T.A., Grootswagers, T., Robinson, A.K., 2019. An introduction to time-resolved decoding analysis for M/EEG.

Cichy, R.M., Pantazis, D., Oliva, A., 2014. Resolving human object recognition in space and time. Nat. Neurosci. 17, 455–462. https://doi.org/10.1038/nn.3635

Clark, A., 2013. Whatever next? Predictive brains, situated agents, and the future of cognitive science. Behav. Brain Sci. 36, 181–204. https://doi.org/10.1017/S0140525X12000477

Dale, A.M., Fischl, B., Sereno, M.I., 1999. Cortical Surface-Based Analysis. Neuroimage 9, 179–194. https://doi.org/10.1006/nimg.1998.0395

Demarchi, G., Sanchez, G., Weisz, N., 2019. Automatic and feature-specific prediction-related neural activity in the human auditory system. Nat. Commun. 10, 1–11. https://doi.org/10.1038/s41467-019-11440-1

Den Ouden, H.E.M., Friston, K.J., Daw, N.D., McIntosh, A.R., Stephan, K.E., 2009. A dual role for prediction error in associative learning. Cereb. Cortex 19, 1175–1185. https://doi.org/10.1093/cercor/bhn161

Dima, D.C., Perry, G., Messaritaki, E., Zhang, J., Singh, K.D., 2018. Spatiotemporal dynamics in human visual cortex rapidly encode the emotional content of faces. Hum. Brain Mapp. 39, 3993–4006. https://doi.org/10.1002/hbm.24226

Dima, D.C., Singh, K.D., 2018. Dynamic representations of faces in the human ventral visual stream link visual features to behaviour.

Fan, R.E., Chang, K.W., Hsieh, C.J., Wang, X.R., Lin, C.J., 2008. LIBLINEAR: A library for large linear classification. J. Mach. Learn. Res. 9, 1871–1874. https://doi.org/10.1145/1390681.1442794

Friston, K.J., Holmes, A.P., Worsley, K.J., Poline J.-P, Frith, C.D., Frackowiak, R.S.J., 1994. Statistical parametric maps in functional imaging: A general linear approach. Hum. Brain Mapp. 2, 189–210. https://doi.org/10.1002/hbm.460020402

Grill-Spector, K., Henson, R., Martin, A., 2006. Repetition and the brain: Neural models of stimulus-specific effects. Trends Cogn. Sci. 10, 14–23.

Grootswagers, T., Wardle, S. G., & Carlson, T. A. (2017). Decoding dynamic brain patterns from evoked responses: A tutorial on multivariate pattern analysis applied to time series neuroimaging data. Journal of cognitive neuroscience, 29(4), 677–697.

Guggenmos, M., Sterzer, P., Cichy, R.M., 2018. Multivariate pattern analysis for MEG: A comparison of dissimilarity measures. Neuroimage 173, 434–447. https://doi.org/10.1016/j.neuroimage.2018.02.044

Hogendoorn, H., Burkitt, A.N., 2018. Predictive coding of visual object position ahead of moving objects revealed by time-resolved EEG decoding. Neuroimage 171, 55–61. https://doi.org/10.1016/j.neuroimage.2017.12.063

JASP Team, 2020. JASP. [Computer software].

John-Saaltink, E.S., Utzerath, C., Kok, P., Lau, H.C., De Lange, F.P., 2015. Expectation suppression in early visual cortex depends on task set. PLoS One 10, 1–14. https://doi.org/10.1371/journal.pone.0131172

King, J.R., Pescetelli, N., Dehaene, S., 2016. Brain Mechanisms Underlying the Brief Maintenance of Seen and Unseen Sensory Information. Neuron 92, 1122–1134. https://doi.org/10.1016/j.neuron.2016.10.051

Kok, P., Mostert, P., De Lange, F.P., 2017. Prior expectations induce prestimulus sensory templates. Proc. Natl. Acad. Sci. U. S. A. 114, 10473–10478. https://doi.org/10.1073/pnas.1705652114

Kok, P., Rahnev, D., Jehee, J.F.M., Lau, H.C., De Lange, F.P., 2012. Attention reverses the effect of prediction in silencing sensory signals. Cereb. Cortex 22, 2197–2206. https://doi.org/10.1093/cercor/bhr310

Larsson, J., Smith, A.T., 2012. FMRI repetition suppression: Neuronal adaptation or stimulus expectation? Cereb. Cortex 22, 567–576. https://doi.org/10.1093/cercor/bhr119

Maunsell, J.H.R., Gibson, J.R., 1992. Visual response latencies in striate cortex of the macaque monkey. J. Neurophysiol. 68, 1332–1344. https://doi.org/10.1152/jn.1992.68.4.1332

Mechelli, A., Price, C.J., Friston, K.J., Ishai, A., 2004. Where bottom-up meets top-down: Neuronal interactions during perception and imagery. Cereb. Cortex 14, 1256–1265. https://doi.org/10.1093/cercor/bhh087

Mumford, D., 1992. On the computational architecture of the neocortex - II The role of cortico-cortical loops. Biol. Cybern. 66, 241–251. https://doi.org/10.1007/BF00198477

Oostenveld, R., Fries, P., Maris, E., Schoffelen, J.M., 2011. FieldTrip: Open source software for advanced analysis of MEG, EEG, and invasive electrophysiological data. Comput. Intell. Neurosci. 2011. https://doi.org/10.1155/2011/156869

Pantazis, D., Fang, M., Qin, S., Mohsenzadeh, Y., Li, Q., Cichy, R.M., 2018. Decoding the orientation of contrast edges from MEG evoked and induced responses. Neuroimage 180, 267–279. https://doi.org/10.1016/j.neuroimage.2017.07.022

Spratling, M.W., 2017. A review of predictive coding algorithms. Brain Cogn. 112, 92–97. https://doi.org/10.1016/j.bandc.2015.11.003

Utzerath, C., St John-Saaltink, E., Buitelaar, J., De Lange, F.P., 2017. Repetition suppression to objects is modulated by stimulus-specific expectations. Sci. Rep. 7, 1–8. https://doi.org/10.1038/s41598-017-09374-z

Veen, B.D. Van Drongelen W. Van Yuchtman, M., Suzuki, A., 1997. Localization of brain electrical activity via linearly constrained minimum variance spatial filtering. IEEE Transactions on. Biomed. Eng. (NY). 44, 867–880.

Walsh, K.S., Mcgovern, D.P., Clark, A., Connell, R.G.O., 2020. Evaluating the neurophysiological evidence for predictive processing as a model of perception 1–27. https://doi.org/10.1111/nyas.14321

